# Internally generated population activity in cortical networks hinders information transmission

**DOI:** 10.1101/2020.02.03.932723

**Authors:** Chengcheng Huang, Alexandre Pouget, Brent Doiron

## Abstract

How neuronal variability impacts neuronal codes is a central question in systems neuroscience, often with complex and model dependent answers. Most population models are parametric, with a tacitly assumed structure of neuronal tuning and population-wide variability. While these models provide key insights, they purposely divorce any mechanistic relationship between trial average and trial variable neuronal activity. By contrast, circuit based models produce activity with response statistics that are reflection of the underlying circuit structure, and thus any relations between trial averaged and trial variable activity are emergent rather than assumed. In this work, we study information transfer in networks of spatially ordered spiking neuron models with strong excitatory and inhibitory interactions, capable of producing rich population-wide neuronal variability. Motivated by work in the visual system we embed a columnar stimulus orientation map in the network and measure the population estimation of an orientated input. We show that the spatial structure of feedforward and recurrent connectivity are critical determinants for population code performance. In particular, when network wiring supports stable firing rate activity then with a sufficiently large number of decoded neurons all available stimulus information is transmitted. However, if the inhibitory projections place network activity in a pattern forming regime then the population-wide dynamics compromise information flow. In total, network connectivity determines both the stimulus tuning as well as internally generated population-wide fluctuations and thereby dictates population code performance in complicated ways where modeling efforts provide essential understanding.

## Introduction

A prominent feature of cortical response to sensory stimuli is that neuronal activity varies significantly across presentation trials^1,2^, even when efforts are taken to control or account for variable animal behavior^3–6^. A component of this variability is coordinated among neurons in a brain area, often leading to shared fluctuations in spiking activity^7–11^. How stimulus processing is affected by this large, population-wide neuronal variability is a longstanding question in both experimental and theoretical neuroscience communities.

Recording from neuronal populations while simultaneously monitoring an animal’s behavior during a structured task offers a glimpse into how neuronal activity supports computation. Correlations between spike counts from pairs of neurons in response to repeated stimulus, often referred to as noise correlations, are modulated by a variety of cognitive factors that are known to affect task performance^4^. For example, noise correlations decrease with animal arousal^12,13^ or task engagement^14^. In the visual pathway noise correlations are decreased when spatial attention is directed into the receptive field of a recorded population^15,16^. In addition, perceptual learning^17,18^ and visual experience^19^ can also result in an attenuation of noise correlations. The common theme in all of these studies is that a reduction in noise correlations co-occurs with cognitive shifts that improve task performance. This supports the often cited idea that shared variability is deleterious to neuronal coding because it cannot be reduced by ensemble averaging^20,21^, and thus it is expected that its reduction will enhance neural coding.

While the above narrative is appealing, it oversimplifies a long debate in the computational neuroscience community^20,22–26^. It is popular (and pragmatic) to restrict analysis to a linear decoding of neuronal response, where performance is often measured using the linear Fisher information between the estimated and actual stimulus^23,27–30^. Linear Fisher information depends on two components: the set of neuronal tuning curves and the population covariance matrix^23,27,28^. Whether noise correlations degrade or improve stimulus coding depends on the relationship between these components. For example, while it is true that correlations between similarly tuned neurons limit Fisher information^21,31^, correlations between dissimilarly tuned neurons can actually increase information compared to an asynchronous population^22,23^. Several more recent studies have built upon this idea and shown how stimulus dependent correlations can also improve linear information transmission^32–34^. Finally, Moreno-Bote and colleagues^35^ showed that information is only limited by one specific type of correlations, termed differential correlations, which align with population activity in the direction defined by the gain of population tuning. In all of these varied modelling studies both tuning curves and covariance structure were assumed and thus had a prescribed relation to one another. In this study we adopt an alternative modelling approach and consider biophysically grounded population models, where cellular spiking dynamics and synaptic wiring are assumed, and population response and its variability are emergent properties of the circuit.

A serious obstacle in using circuit models to study neuronal coding is the lack of complete mechanistic theory underlying population-wide variability^36^. An often used framework assumes neuronal fluctuations are inherited from external sources, and any circuit wiring filters and propagate this variability throughout the population^37–40^. Along these lines recent modelling work in a feedforward circuit model of primary visual cortex shows that when such inherited variability originates from a noisy stimulus then population-wide correlations which limit information transfer naturally develop throughout the circuit^41^. Neuronal variability can also emerge through recurrent interactions within the network, when synaptic weights are large and excitation is dynamically balanced by recurrent inhibition^1,42,43^. However, these networks famously support an asynchronous state^44^, making them at the surface ill-suited to probe how internally generated population-wide variability affects neuronal processing. Recent extensions of balanced networks which include structured wiring has provided an understanding of how shared variability can emerges through internal mechanics^45–52^. These networks are a useful modelling framework to probe how circuit produced shared variability impacts population coding.

In this work, we study information transmission in spatially ordered spiking neuron networks where a balance between excitation and inhibition produce population-wide variability. Modelling simple visual inputs to cortex with a columnar organization of stimulus orientation preference allows a study of how neuronal stimulus tuning and associated population variability are shaped by circuit wiring and modulatory inputs. We found that narrow feedforward projections increase information in the output layer for finite number of decoding neurons, while the recurrent projection width with spatially balanced excitation and inhibition has little effect. Interestingly, the total information is preserved when the network is in a regime where firing rates are stable. In contrast, when the network exhibits coherent internal dynamics through a pattern forming instability in firing rates, the information transfer is largely reduced. Our work thus begins to connect the emerging theory of how neuronal circuits produce trial-variable activity with ongoing theories of how population-wide variability affect stimulus representation.

## Results

Our study measures the propagation of stimulus information in a two-layer network of spatially ordered neurons. As motivation our model aims to capture layer (L)4 and L2/3 neurons in primary visual cortex (V1) (Fig. 1A, see Methods), however our model is sufficiently general to capture an arbitrary layered network. We model L4 neuronal spiking activity as a Poisson process with Gabor receptive fields defining the stimulus to firing rate transfer. The orientation preference of L4 neurons are determined from a superimposed pinwheel orientation map (Fig. 1A, bottom right; see Ref.^53^). The Gabor visual images are corrupted by spatially independent noise obeying a temporal Ornstein-Uhlenbeck process; this provides a bound on the stimulus information entering our network. L2/3 is modeled as a recurrently connected network of both excitatory and inhibitory spiking neuron models. The connection probability of all recurrent projections within L2/3 and the feedforward projections from L4 to L2/3 decay with distance with spatial widths *α*_rec_ and *α*_ffwd_, respectively (see Methods and Ref.^47^ for details).

**Figure 1:**
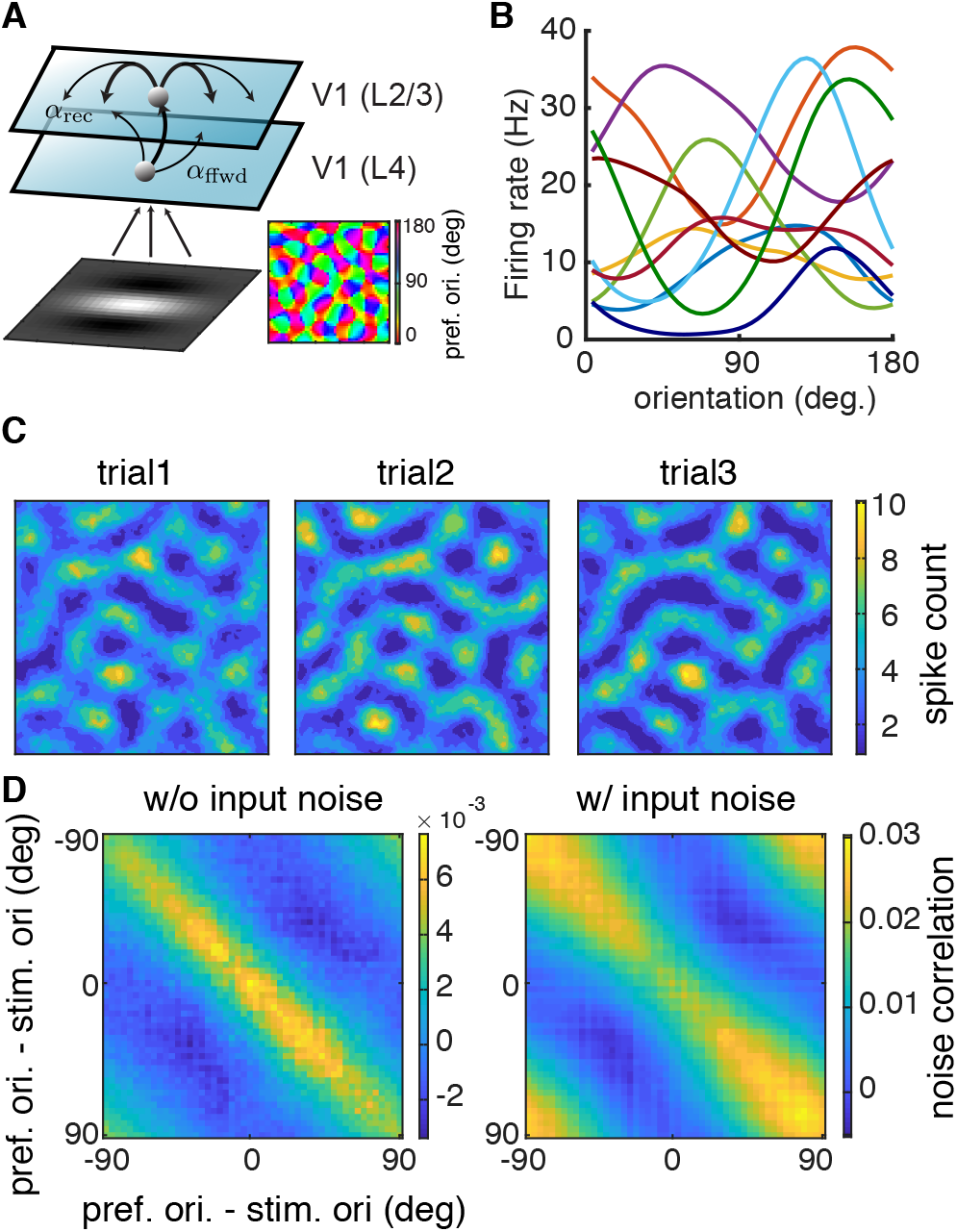
Spatially ordered balanced networks generate heterogeneous tuning curves and structured trial-to-trial variability. **A,** Model schematic of a two-layer network of spatially ordered spiking neurons modeling Layer 4 neurons and Layer 2/3 neurons, respectively, from visual area V1. The visual input to the model is a Gabor image with orientation *θ*. The L4 network consists of Poisson units with Gabor receptive fields. The orientation preferences of L4 neurons are assigned according to a pinwheel orientation map (bottom right). The L2/3 network consists of both excitatory and inhibitory neurons modeled with integrate-and-fire dynamics, all arranged on a unit square. The spatial widths of the feedforward projections from L4 to L2/3 and the recurrent projections within L2/3 are denoted as *α*_ffwd_ and *α*_rec_, respectively. **B,** The L2/3 neurons have heterogeneous orientation tuning curves. Ten examples of tuning curves are shown with different color representing different neurons (smoothed with a Gaussian kernel of width 9°). **C,** The model internally generates trial-to-trial variability. Three trials of network spike counts (200 ms time window) are shown from a network with *α*_ffwd_ = 0.05 and *α*_rec_ = 0.1. Images are smoothed with a Gaussian kernel of width 0.01. D, Noise correlation matrix with neurons ordered by their preferred stimulus orientation. Responses were simulated for a Gabor input with orientation at *θ* = 0° without (left) and with stimulus noise (right).

Networks with large, balanced excitatory and inhibitory connections are an attractive model framework for cortical activity because they capture several aspects of reported *in vivo* population response. When large synaptic connections are paired with random network wiring they naturally produce significant heterogeneity in spiking activities across the network^42,54,55^. Indeed, in our spatially ordered network the L2/3 neurons have very heterogeneous tuning curves with various widths and magnitudes (Fig. 1B). This produces a large spread of orientation selectivity across the population (Figs. 1B, 2B), capturing the broad heterogeneity of orientation tuning reported in primary visual cortex^56^.

**Figure 2:**
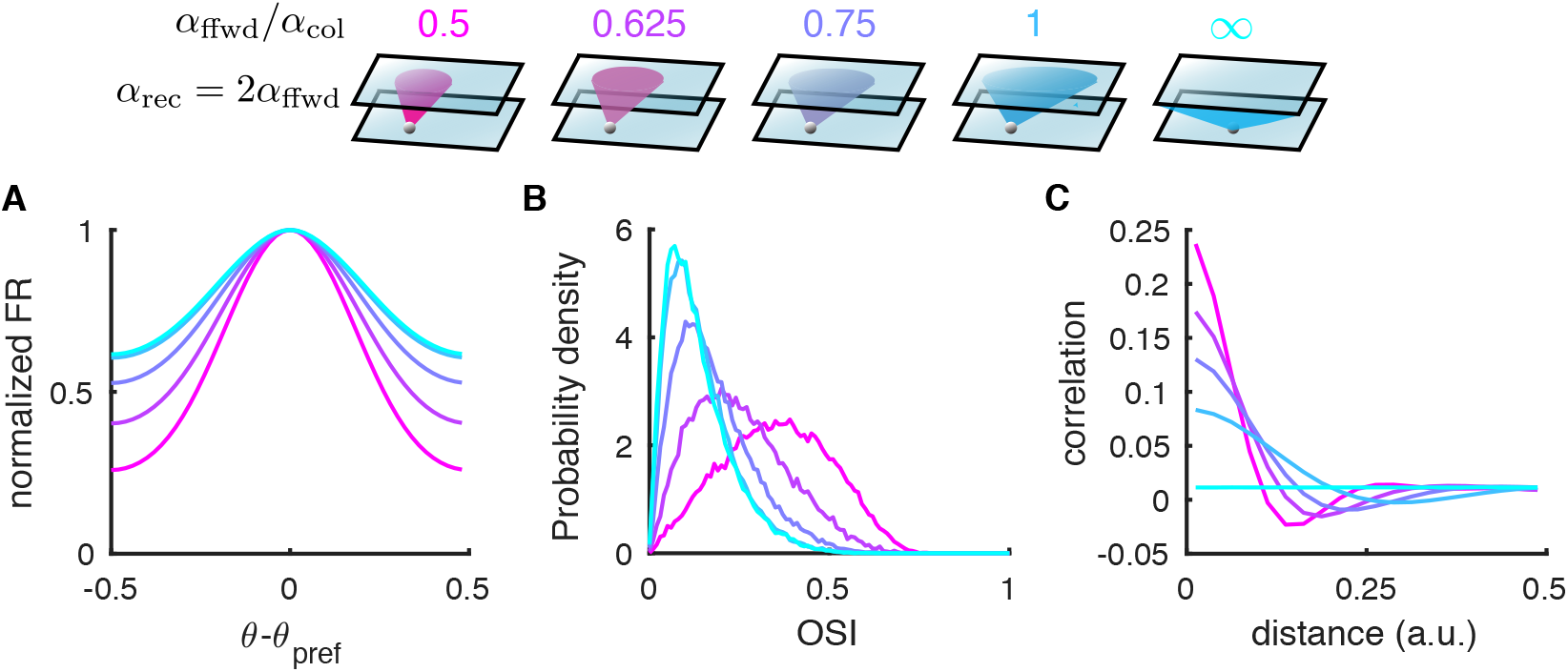
Both tuning curves and noise correlations change with the feedforward projection width. **A,** The average tuning curve, normalized and centered at the preferred orientation, broadens as the feedforward projection width, *α*_ffwd_, increases (color scheme shown on top panel). The recurrent width is fixed to be 2*α*_ffwd_. The columnar radius, *α*_col_, is 0.1. The case of *α*_ffwd_ = ∞ means the connection probability is independent of distance. **B,** The probability density distributions of orientation selectivity indexes (OSI, Eq. 6) of the L2/3 excitatory neurons. **C,** Pairwise correlations as a function of distance between neuron pairs from the L2/3 excitatory population. The magnitude of pairwise correlations decreases as *α*_ffwd_ increases.

Balanced networks produce significant dynamic and trial-to-trial spiking variability through internal mechanisms^42,57^. While balanced networks with disordered connectivity produce asynchronous activity^44^, networks with structured wiring can produce correlated variability^45,48^, that under certain conditions can be population-wide^47^. These past models were concerned with the mechanics of neuronal variability and did not model a spatially distributed stimulus to drive network response. In our model when L4 columnar stimulus structure is enforced then a broad L2/3 columnar activation is recruited accompanied by significant population-wide trial-to-trial variability (Fig. 1C). Despite the lack of feature-based coupling (i.e connectivity was determined only by the spatial distance between neurons), there is a clear positive relation between pairwise signal and noise correlations (Fig. 1D), matching diverse in vivo datasets^15,17,40,58,59^. This occurs for two reasons. First, the stimulus has a spatial columnar profile, so any spatial wiring does partially overlap with the signal profile. Second, it is well known that spike count correlations increase with the firing rates of the neuron pair^60,61^, so that when neuron pairs with similar stimulus preference are co-activated then any significant trial-to-trial covariability in their synaptic inputs is better expressed in their output spiking. In our model, the relation between signal and noise correlations occurs in response to Gabor images which are noiseless (Fig. 1D, left) or those contaminated by sensory noise (Fig. 1D, right). The latter case naturally provides feedforward noise correlations^41^ with a columnar spatial scale. These externally imposed correlations are reduced for neuron pairs with a stimulus preference that match the driving stimulus, in agreement with past models where correlations are inherited from outside the circuit^40^. In total, our spiking network captures many of the trial-averaged and trial-variable aspects of real cortical population response.

We focus on the network’s ability to discriminate two Gabor images with similar orientations, a paradigm commonly used in experiments^62^. Gabor images with orientation *θ* are presented to the L4 neurons repetitively with an ON interval of 200 ms and an OFF interval of 300 ms (see Methods). In what follows we consider a decoder that has access to *N* model L2/3 pyramidal neurons (randomly chosen from the 4×10^4^ L2/3 neurons). We simulate the spiking activities of L2/3 neurons and collect spike counts from the observed population, **n** = [*n*_1_,…, *n_N_*], during each stimulus presentation (ON interval). The linear Fisher information about *θ* available from **n** is defined as:

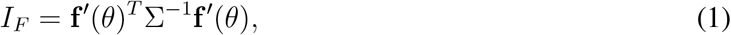

where **f** is the expectation of **n** conditioned on *θ* (population tuning curve),′ denotes differentiation with respect to **θ**, and Σ is the covariance matrix of **n** (noise covariance matrix). Fisher information is a useful metric when considering the estimation of orientation *θ* from **n**. Let 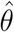 be the optimal linear estimator of the *θ* for a given Gabor image, then we have that 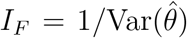^63^. Put more simply, *I_F_* measures the accuracy of the optimal linear estimator of orientation. In practice we measure the linear Fisher information from the spike counts of the L2/3 excitatory neurons using a bias-corrected estimation (see Methods Eq. (7) in Ref.^64^).

The spatial scales of connectivity in our network are key determinants of overall population spiking activity in unstimulated conditions^45–47^. In the next sections we explore how the spatial scales of the feedforward L4 to L2/3 connectivity (*α*_ffwd_; Fig. 1A) and those of the L2/3 to L2/3 recurrent projections (*α*_rec_; Fig. 1A) influence information transfer about stimulus *θ* within the network.

### Narrow feedforward projections increase information saturation rate

Linear Fisher information (Eq. 1) depends on two components: the trial-averaged response gain to stimulus orientation (**f**′) and the trial-to-trial covariance matrix Σ. As we will show, the joint influence of *α*_ffwd_ on gain **f**′ and covariance Σ sets a conflict between how *α*_ffwd_ should ultimately affect *I_F_*. A similar situation can easily exist in models where the form of **f**′ and Σ are assumed^21,23,31–33^. However, in many of these cases the justification for how **f**′ and Σ are related is heuristic, and will depend on the specific parametrization of the response model. While our circuit model also has parameters, such as *α*_ffwd_, there is a mechanistic understanding for how these control population responses, and consequently for how they relate **f**′ and Σ to one another.

On one hand, the population averaged tuning curve broadens as *α*_ffwd_ increases, resulting in reduced orientation gain **f**′ (Fig. 2A). In agreement, the orientation selectivity index (OSI, Eq. 6) of individual neurons decreases with *α*_ffwd_ (Fig. 2B). This reduction in orientation tuning is due to broader spatial filtering of tuned inputs from L4, where the tuning preferences of neurons are spatially clustered in a columnar organization. In particular, when the connection probabilities of all projections are uniform in space (*α*_ffwd_ → ∞), the columnar structure is not preserved in L2/3 and neurons are least tuned. Hence, considering these changes in tuning selectivity alone predict that the IF about θ from L2/3 neurons decoding would decrease with *α*_ffwd_.

On the other hand, the noise correlations between nearby neurons decrease with increasing *α*_ffwd_ (Fig. 2C). Rosenbaum and colleagues^46^ showed that for spatially distributed balanced networks where *α*_ffwd_ < *α*_rec_ an asynchronous solution does not exist. Rather, network activity is organized so that nearby cell pairs have positive noise correlations while more distant cell pairs have negative correlations (Fig. 2C, pink curves). Previous work on population coding has suggested that positive correlation between similarly tuned neurons can limit information while negative correlation between oppositely tuned neurons can benefit coding^20–22^. Since the tuning preferences of neurons are spatially clustered with a pinwheel orientation map, nearby neurons are likely to share similar tuning. Therefore, in contrast to the effect of the tuning selectivity which would suggest that IF could increase with *α*_ffwd_, the overall reduction in the spatial structure of pairwise correlations implies that IF could increase with *α*_ffwd_.

We divide our analysis of how *I_F_* depends on *α*_ffwd_ into two cases: first, when the decoder is restricted to a small (*N* ~ 10^2^) populations of neurons, and second for decoders that have access to large (*N* ~ 10^4^) populations.

For decoders restricted to small populations the combined effects of reduced tuning selectivity and correlations with larger *α*_ffwd_, results in the linear Fisher information being largely reduced as *α*_ffwd_ increases (Fig. 3A). Consistently, the neural thresholds of single neurons, measured as the ratio of the standard deviation of the spike counts and the derivative of the tuning curve function 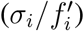, also increase with *α*_ffwd_ (Fig. 3B). In total, the population code is less informative with larger *α*_ffwd_, in line with the single neuron reduction in orientation selectivity (Fig. 2B), despite the associated decrease in pairwise noise correlations (Fig. 2C).

**Figure 3:**
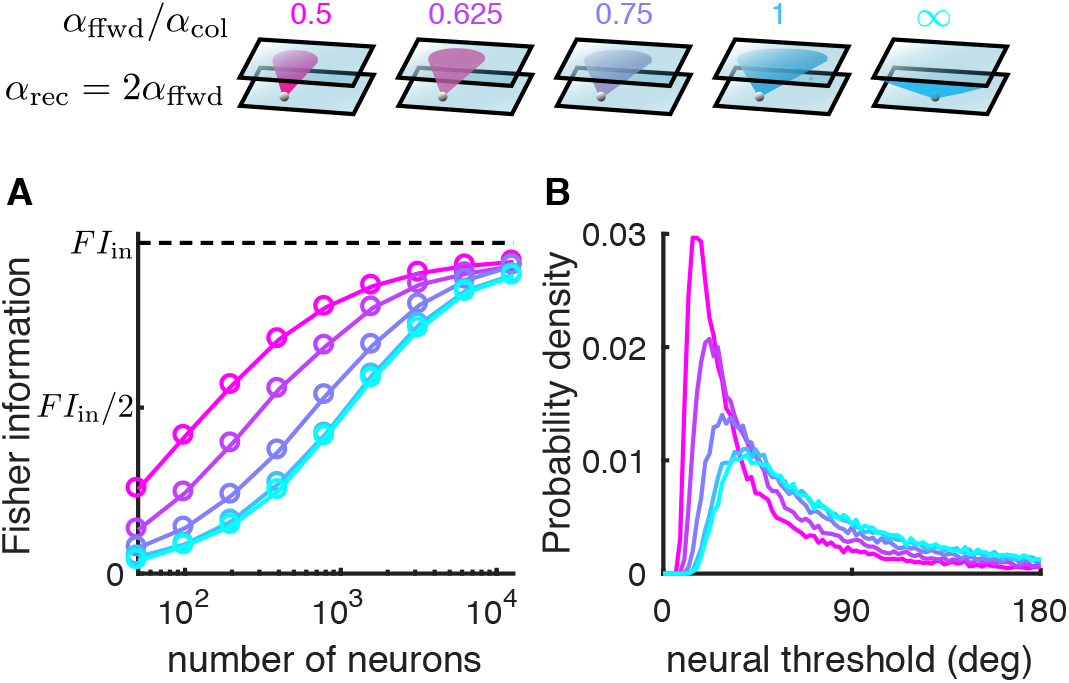
The linear Fisher information saturates faster with smaller feedforward projection width. **A,** The linear Fisher information increases with the number of neurons sampled from the excitatory population in L2/3 (color scheme in top panel). With large number of neurons, the linear Fisher information saturates close to the input information bound from the L4 network (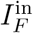, dashed line). The input noise on the Gabor images is chosen such that 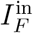 corresponds to a discrimination threshold of about 1.8 degrees, which is consistent with psyphophysical experiments^65^ (see Methods). **B,** The probability density distributions of neural thresholds 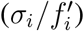 of the L2/3 excitatory neurons.

The noise in the Gabor image creates shared fluctuations in the L2/3 neurons that cannot be distinguished from signal, limiting the available information about *θ* that is possible to be decoded (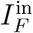; Fig. 3A dashed line, see Methods). For decoders that have access to large populations the linear Fisher information of the L2/3 neurons saturates to a level close to 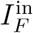, regardless of the feedforward projection width (Fig. 3A). It suggests that the network is very efficient at transmitting information. In particular, even networks with spatially disordered connections (*α*_ffwd_ → ∞), where neurons are very weakly tuned, still contain most of the information from the input layer.

Moreno-Bote and colleagues^35^ explicitly make the observation that while networks may show significant noise correlations, it is the shared fluctuations that align with the direction of response gain that limit information. In particular, they consider the decomposition of the covariance Σ = Σ_0_ + *ϵ***f′f′^*T*^**, with *ϵ* measuring the shared variance along the coding direction (**f**′). They show that for decoders that have access to a large number of neurons then *I_F_* ~ 1/*ϵ*. For our network it is not (to our knowledge) possible to formally decompose Σ into Σ_0_ + *ϵ***f′f′^*T*^**, meaning we do not have a way of estimating how network interactions contribute to *ϵ*. However, the fact that the information eventually saturates close to 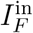 for all values of *α*_ffwd_ indicates that *α*_ffwd_ does not affect the value of *ϵ*. This is consistent with our network being in a regime where the origin of information limiting correlations stems from strictly external fluctuations, as has been previously studied^41^.

### Recurrent connections with spatially balanced excitation and inhibition have small effects on information transfer

Next we study how the spatial scales of recurrent connections (*α*_rec_) affect information transmission. We first consider networks with the same spatial scale of recurrent excitatory and inhibitory projections. As remarked in the previous section, the relative spatial scale between recurrent and feedforward projections has a large impact on the spatial structure of pairwise correlations (Fig. 2C, Ref.^46^). For fixed *α*_ffwd_ then as *α*_rec_ increases orientation selectivity increases (Fig. 4C), while pairwise noise correlations become spatially structured (Fig. 4D).

**Figure 4:**
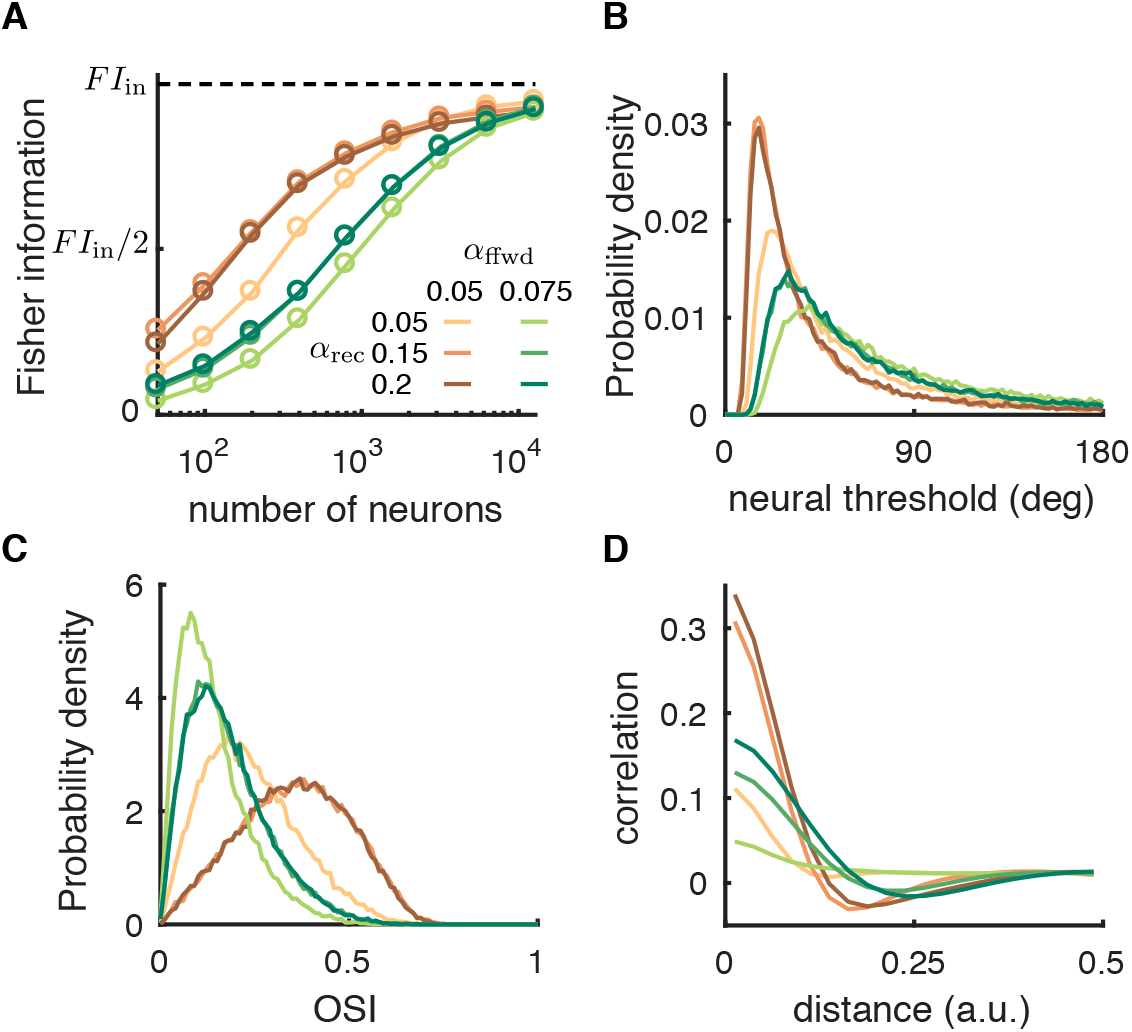
The recurrent projection width has small effects on the linear Fisher information. **A,** The linear Fisher information is slightly lower when the recurrent projection width (*α*_rec_) is narrower than the feedforward width (*α*_ffwd_) (light shade). When *α*_rec_ is broader than *α*_ffwd_, the saturation curve of the Fisher information overlaps (darker shades). Examples of two *α*_ffwd_ values are shown (red: *α*_ffwd_ = 0.05; green: *α*_ffwd_ = 0.075). B, The probability density distributions of neural thresholds 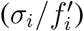 of the L2/3 excitatory neurons. **C,** The probability density distributions of orientation selectivity indexes of the L2/3 excitatory neurons. **D,** Pairwise correlation as a function of the distance between a neuron pair.

For decoders that consider only small populations (*N* ~ 10^2^ to 10^3^ neurons) we see a clear difference between how *α*_rec_ and *α*_ffwd_ affect IF. Specifically, IF is relatively insensitive to *α*_rec_ (the brown curves scanning over *α*_rec_ for *α*_ffwd_ = 0.05 are clustered in Fig. 4A; similar for the green curves with *α*_ffwd_ = 0.075). Indeed, both the distributions of neural thresholds and OSI show similar ordering as the Fisher information (Fig. 4B,C). Further, the spatial structures of pairwise correlations do not contribute to a difference in Fisher information (Fig. 4D), again suggestive that the population correlations are not information limiting. In agreement, for decoders with access to large populations *I_F_* saturates to 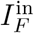. Overall, for spatially balanced recurrent excitatory and inhibitory projections the specific scale of *α*_rec_ is of little importance when considering information transfer.

### Broad inhibitory projections reduce information

Our cortical network can exist in one of two regimes. The preceding sections consider a regime where circuit parameters are such that the firing rate dynamics across the network are stable, despite pairwise correlations being structured. The second regime is where the firing rates are no longer dynamically stable, and large rate fluctuations are produced through internal interactions (as we will formalize below). In past work^47^ we showed how model networks in this second regime produced the low-dimensional, population-wide shared variability characteristic of a variety of cortical areas^9–11,47,66–69^. In this section we consider how networks in this second regime transfer information across layers.

Dynamical networks with spatially ordered coupling can produce rich spatio-temporal patterns of activity^70^. To explore how the network circuitry in our cortical model determines its pattern forming dynamics we first implement two simplifications. First, we restrict our analysis to an associated firing rate model^71,72^, sharing the same spatial connectivity as our network of model spiking neurons (see Methods). This greatly simplifies any linear stability analysis, and it provides qualitatively similar macroscopic network firing rates when spiking neuron models are in a fluctuation driven regime^73,74^. Second, we analyze the dynamics of the network when driven by spatially uniform inputs. This provides a spatial translational symmetry in the network needed for Fourier analysis.

A stable, spatially uniform solution of the firing rate model corresponds to an asynchronous state in the network of spiking neuron models. We linearize the firing rate network dynamics around this uniform solution and obtain a set of eigenvalues and associated eigenmodes. This eigenstructure governs the linearized firing rate dynamics near the uniform solution. Each eigenmode has a wavenumber indicating its spatial scale across the network; the zero wavenumber eigenmode describes a spatially uniform solution, while higher wavenumber eigenmodes contribute to spatially structured solutions (at the spatial period of the wavenumber). The stability of the solution at each eigenmode is given by the sign of the real component of the associated eigenvalue: negative (positive) eigenvalues imply dynamics about that eigenmode are stable (unstable). If all eigenmodes are stable then we say that the solution is stable.

Networks with spatially balanced recurrent excitatory and inhibitory projections (*σ_e_* = *σ_i_*) and strong static inputs to the inhibitory neurons (*μ_i_*) have a stable spatially uniform solution (Figs. 5A, grey region; 5B, orange curves). In this regime, the network dynamics is approximately linear with weak perturbations, meaning that the network fluctuations are linearly related with input fluctuations. Since linear Fisher information is conserved under an invertible linear transformation (see Discussion), a network in a stable regime preserves almost all of its input information. Indeed, as shown in previous sections, *I_F_* converges to 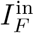 for large *N* decoders, and this convergence is independent of the connectivity scales of both the feedforward and the recurrent pathways (Fig. 3A, Fig. 4A).

**Figure 5:**
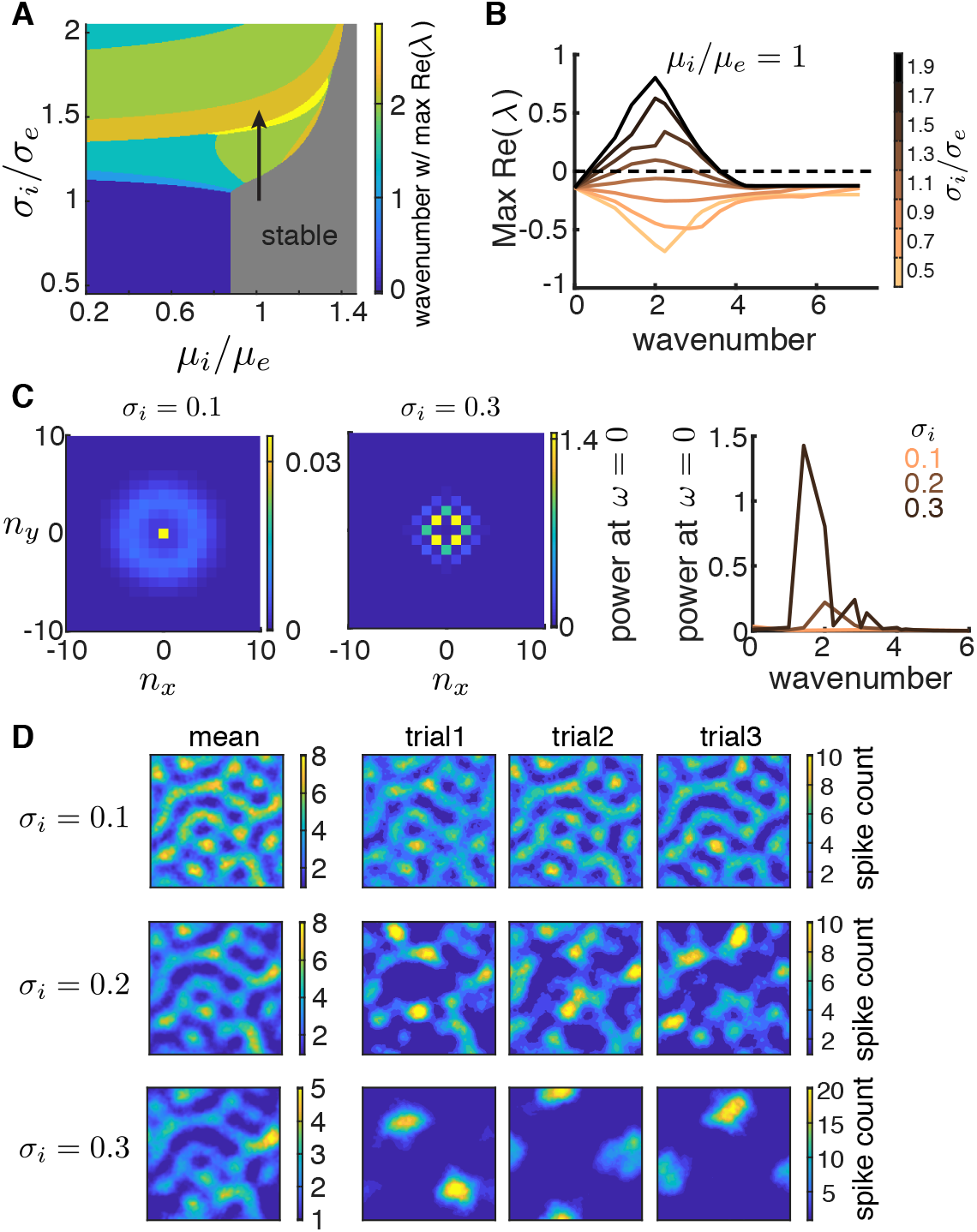
Networks with broad inhibitory projections exhibit pattern forming dynamics. **A,** Bifurcation diagram of a firing rate model as a function of the inhibitory projection width *σ_i_* and the depolarization current to the inhibitory neurons (*μ_i_*). The excitatory projection width and the drive to the excitatory neurons are fixed at *σ_e_* = 0.1 and *μ_e_* = 0.5, respectively. Color represents the wavenumber whose eigenvalue has the largest positive real part and the gray region is marked stable since all eigenvalues have a negative real part. **B,** The maximum real part of eigenvalues as a function of wavenumber for increasing *σ_i_* (*σ_e_* = 0.1), when *μ_i_ = μ_e_*. Nonzero wavenumbers lose stability as *σ_i_* increases, indicating that the network will exhibit firing rate dynamics with spatial scales of the unstable eigenmodes of the uniform solution. **C,** The power spectrum at temporal frequency *ω* = 0 for different spatial Fourier modes (*n_x_, n_y_*) of the spontaneous spiking activity from the spiking neuron networks with *σ_i_* = 0.1 (left, the nonzero spatial frequency power peaks around wavenumber 3.6.) and *σ_i_* = 0.3 (middle). Right: The average power at *ω* = 0 across wavenumbers 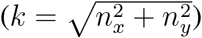. **D,** Activities of spiking neuron networks with different *σ_i_* when driven by a Gabor image (right: three trials of spike counts within 200 ms window; left: mean spike counts). Images are smoothed with a Gaussian kernel of width 0.01. Other parameters for the spiking neuron networks (panels C, D) are *σ_e_* = 0.1, *α*_ffwd_ = 0.05, and feedforward strengths *J_eF_* = 240 mV and *J_iF_* = 400 mV.

When the excitatory and inhibitory projections differ in spatial scale, recurrent connectivity can have a large impact on network dynamics. As the inhibitory projection width (*σ_i_*) increases the uniform solution in the spatial firing rate model loses stability at a band of non-zero wavenumbers (Fig. 5A,B). This is via a Turing-Hopf bifurcation^70^, a pattern-forming transition common to spatially distributed neuronal network models^71,72,75^. The unstable eigenmodes that emerge from the bifurcation contribute to population dynamics confined to the spatial scale of their wavenumbers. Such pattern forming transitions are also observed in the spontaneous activity of spiking neuron networks^45,47,51,76,77^. To quantify the spatial scales of population activity we compute the spatial power of population spiking (at zero temporal frequency) in the spontaneous state. When *σ_i_* equals *σ_e_* the power spectrum is small (Fig. 5C, right, orange curve) showing a band at a wavenumber of approximately 3.6 (Fig. 5C, left), reflecting the spatial scale of the feedforward projections (see Supp Fig 3 in Ref.^46^). As *σ_i_* increases, there is a large increase in power at some non-zero wavenumbers (Fig. 5C), reflecting large, internally generated fluctuations at non-zero spatial frequencies.

When the spiking network is driven by a Gabor image, the internal dynamics of the network interacts with the spatial patterns of the feedforward inputs. In the stable regime, where excitation is spatially balanced with inhibition (*σ_i_ = σ_e_*), the spatial patterns of the network activity are similar across trials and are inherited from the input spatial scale (Fig. 5D, first row). As *σ_i_* increases, the internal pattern forming dynamics of the network become dominant over the feedforward input patterns (Fig. 5D, top to bottom rows). The broad inhibition suppresses significant activity, resulting in only a few stimulus evoked areas compared to the spatially balanced case. The locations of the evoked areas vary from trial to trial, hence introducing excessive trial-to-trial variability in the population activity patterns. The mean population activity pattern across trials also degrades as *σ_i_* increases (Fig. 5D, left). In total, this suggests that there will be information loss in networks with larger *σ_i_*.

Indeed, the Fisher information *I_F_* about stimulus *θ* is markedly reduced when the network loses stability through broader inhibitory projections (Fig. 6A). The orientation selectivity of neurons only changes slightly with *σ_i_* (Fig. 6B), which means that a reduction in tuning selectivity cannot be the primary reason for the decrease in *I_F_*. However, the magnitude of pairwise correlations increases drastically as *σ_i_* increases. Due to the internal pattern forming dynamics in networks with large *σ_i_*, nearby neurons are strongly correlated while neuron pairs that are separated by half the distance between active zones are strongly anti-correlated. Consider the decomposition of network covariance 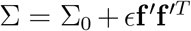 (see Ref.^35^), recalling that in the limit of large decoded populations (*N*) we have that the Fisher information *I_F_* ~ 1/*ϵ*, so that *ϵ* measures the strength of information limiting correlations. In the stable regime *I_F_* converges to the input information 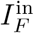 (Fig. 6A, orange curve), implying that all information limiting correlations are inherited through the feedforward pathway (as reported earlier in Figs. 3 and 4). However, for broader inhibition it appears that for large *N* the information *I_F_* saturates to a level that is below 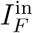. This argues that a component of the internally generated network covariability actually limits information transfer (i.e contributes to *ϵ*). While our simulations give strong support to this conclusion we admit that since it is computationally prohibitive to consider N beyond 10^4^ neurons we cannot verify how *I_F_* truly converges. In total, the pattern-forming dynamics in networks with broad inhibitory projections largely reduce the transmitted information.

**Figure 6:**
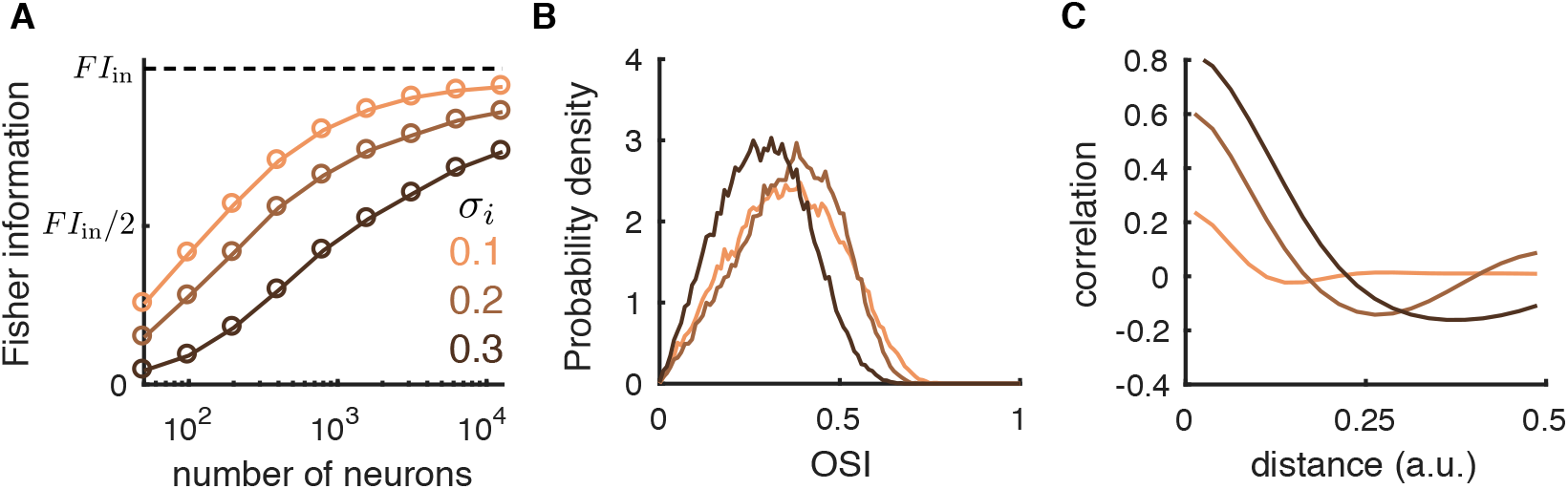
Broad inhibitory projections reduce information flow. **A,** The linear Fisher information from the L2/3 excitatory neurons is largely reduced as the inhibitory projection width (*σ_i_*) increases. **B,** The probability density distributions of orientation selectivity indexes of the excitatory neurons from L2/3. **C,** Pairwise correlations as a function of distance between a neuron pair in L2/3.

### Depolarizing inhibitory neurons improves information flow

Even when excitatory and inhibitory projections are spatially matched a network can nevertheless have an unstable uniform solution. This occurs when the static input to the inhibitory neurons (*μ_i_*) is low (Fig. 7A, blue region) and the firing rate network is unstable through the eigenmode with zero wavenumber (Fig. 7B). In this case the stability of the uniform solution is mediated by a Hopf bifurcation, through which bulk population-wide firing dynamics are produced (Fig. 7A, transition from blue to grey regions), as opposed to the spatially confined dynamics produced via a Turing instability (Fig. 5). The spiking network shows a similar instability; when the static current to the inhibitory neurons is small the spontaneous activities of spiking neuron networks exhibit large power at zero wave number, indicating large magnitude global fluctuations (Fig. 7C). Further, the global fluctuations in the spontaneous activities of the spiking neuron network are largely suppressed with more input to the inhibitory neurons (Fig. 7C). In response to a stimulus θ the population activity patterns show large fluctuations in the overall spiking activity level when μ¿ is small (Fig. 7D). However, unlike the case for broad inhibition, since the instability is at zero wavenumber then no spatially patterned internal dynamics occur that would compete with the stimulus evoked spatial patterns. Consequently, the spatial patterns of the evoked activities are similar across trials (Fig. 7D).

**Figure 7:**
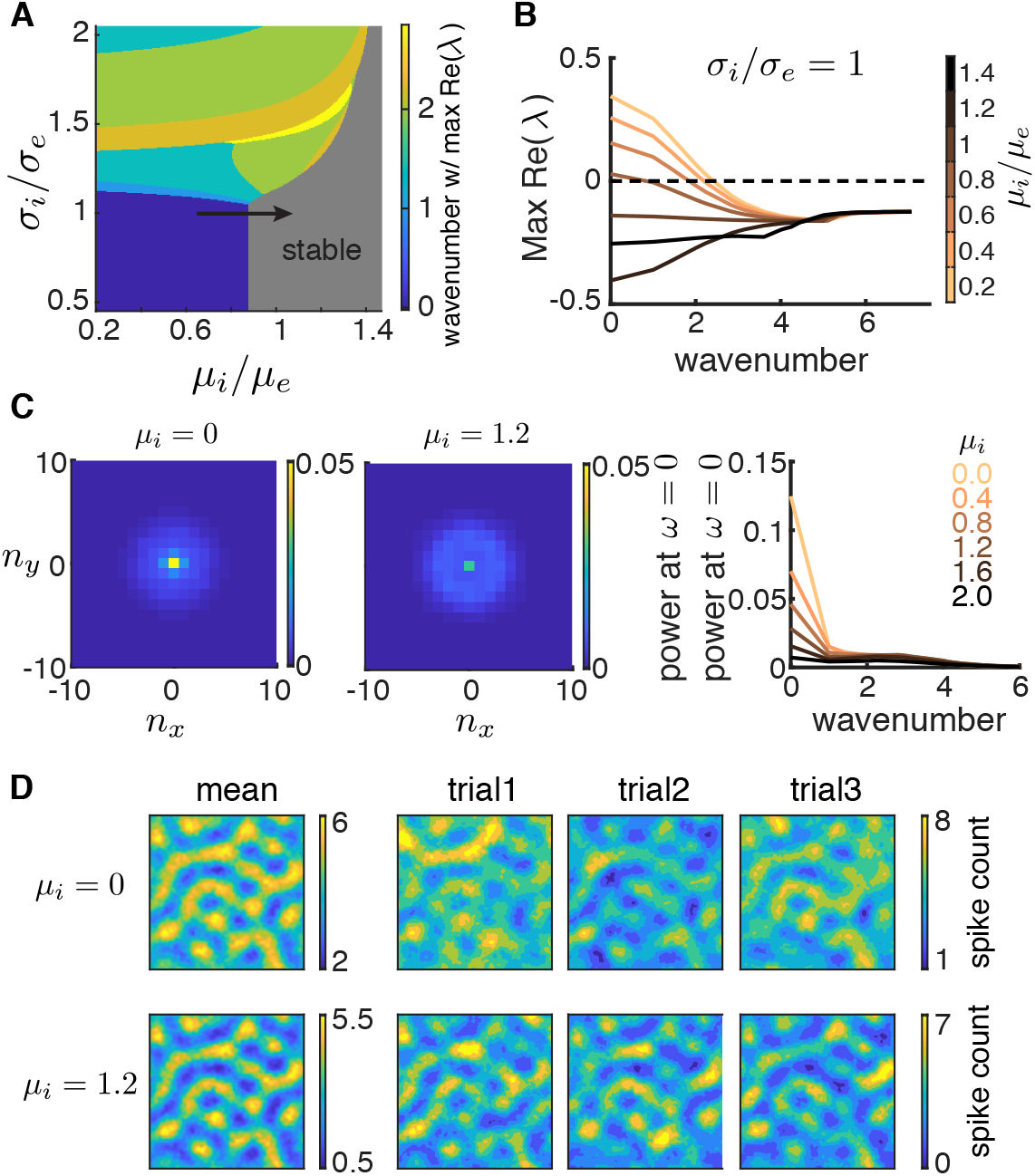
Lack of inhibitory drive gives rise to network-wide fluctuations. **A,** Bifurcation diagram of a firing rate model as a function of the inhibitory projection width σi and the depolarization current to the inhibitory neurons. Same as Fig. 5A. **B,** The real part of eigenvalues as a function of wave number for increasing *μ_i_* (*μ_e_* = 0.5) when *σ_i_ = σ_e_*. The zero wave number loses stability when *μ_i_* is small, indicating that the network will exhibit network-wide nonlinear dynamics. **C,** The power spectrum at temporal frequency *ω* = 0 for different spatial Fourier modes (*n_x_, n_y_*) of the spontaneous spiking activity from the spiking neuron networks with *μ_i_* = 0 (left) and *μ_i_* = 1.2 (middle). Right: The average power at *ω* = 0 across wave number 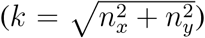. **D,** Activities of spiking neuron networks with different *μ_i_* when driven by a Gabor image (right: three trials of spike counts within 200 ms window; left: mean spike counts). Images are smoothed with a Gaussian kernel of width 0.01. Other parameters for the spiking neuron networks (panels C, D) are *σ_e_ = σ_i_* = 0.1, *α*_ffwd_ = 0.05, *J*_eF_ = 140 mV, *J*_iF_ = 0 mV, and the tonic current to excitatory neurons is *μ_e_* = 0.

The Fisher information about *θ* increases with *μ_i_* as the network becomes more stable (Fig. 8A). Once the network is in the stable regime, a further increase in *μ_i_* has little effect on the *I_F_* (Fig. 8A, *μ_i_* = 0.8 compared *μ_i_* = 1.2). To understand this we again consider how response gain and population covariability are affected by *μ_i_*. First, as *μ_i_* increases the tuning curves of the excitatory neurons are sharpened (Fig. 8B). Second, since the unstable dynamics correlate the whole network, they give rise to positive correlations between neurons across long distances. As *μ_i_* increases these population-wide fluctuations are suppressed and overall neurons are less correlated (Fig. 8C). Indeed, this combination could in principle conspire to produce the increase in information. However, we note that the impact of global fluctuations upon information transfer is much less than that of the spatially confined fluctuations induced by broad inhibition (compare Figs. 6A and 8A).

**Figure 8:**
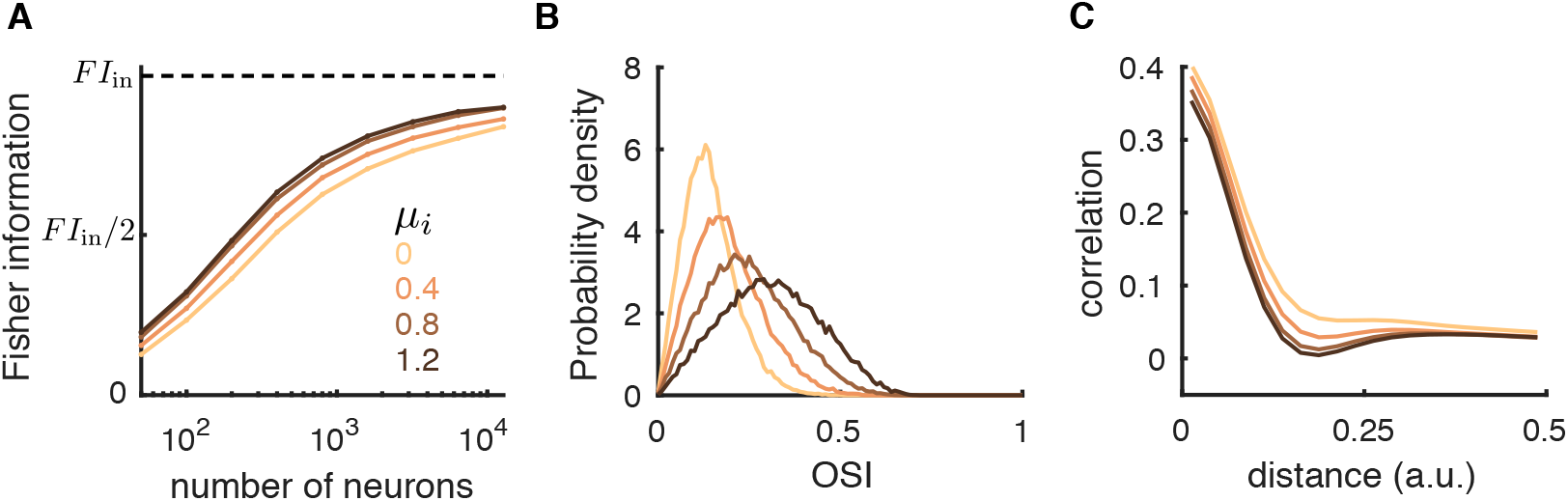
Depolarizing inhibitory neurons improves information flow. **A,** The linear Fisher information increases with depolarizing the inhibitory neurons (*μ_i_*). The saturation curves of the linear Fisher information overlap for *μ_i_* ≥ 0.8. **B,** The probability density distributions of orientation selectivity indexes of the L2/3 excitatory neurons. **C,** Pairwise correlations as a function of the distance between neuron pairs from the L2/3 excitatory population. The feedforward strengths are *J*_eF_ = 140 mV and *J*_iF_ = 0 mV, and the tonic current to excitatory neurons is *μ_e_* = 0.

The small improvement in information transfer between two layers can be compounded during multi-layer processing. We consider a four-layer network model with the same spatial wiring (Fig. 9A). The first and the second layer are the same as the L4 and L2/3 networks, respectively, in previous model. We compare the two networks with different inhibitory biases (*μ_i_*) applied to the neurons in Layers 2, 3, and 4. The feedforward strength was adjusted such that the firing rates were similar across layers 2 to 4 (Supp Fig. S1C), as well as between the two conditions of *μ_i_*. As expected the information about θ decreases as it propagates to higher-order layers (Fig. 9B,C), this is simply an expression of the data-processing inequality^78^. This suggests that information becomes more diffused in higher-order layers (Fig. 9B,C); indeed, the pairwise correlations are higher and the thresholds of single neurons are larger in higher-order layers (Supp Fig. S1A,B). However, the information deteriorates much faster across layers in networks with smaller *μ_i_* (Fig. 9B,C), not only due to the compounding effect, but also because of the increase in synchrony in spike trains among neurons for small *μ_i_* (Supp Fig. S1D). Since the inhibition is insufficient to cancel correlations at each layer, the synchrony builds up as signal propagates to higher-order layers^79,80^. With stronger input to the inhibitory neurons, excitation is more balanced by inhibition, leading to stable asychronous dynamics (Supp Fig. S1D). In total, the improvement in the Fisher information by increasing *μ_i_* dramatically increases with the number of processing layers (Fig. 9D).

**Figure 9.**
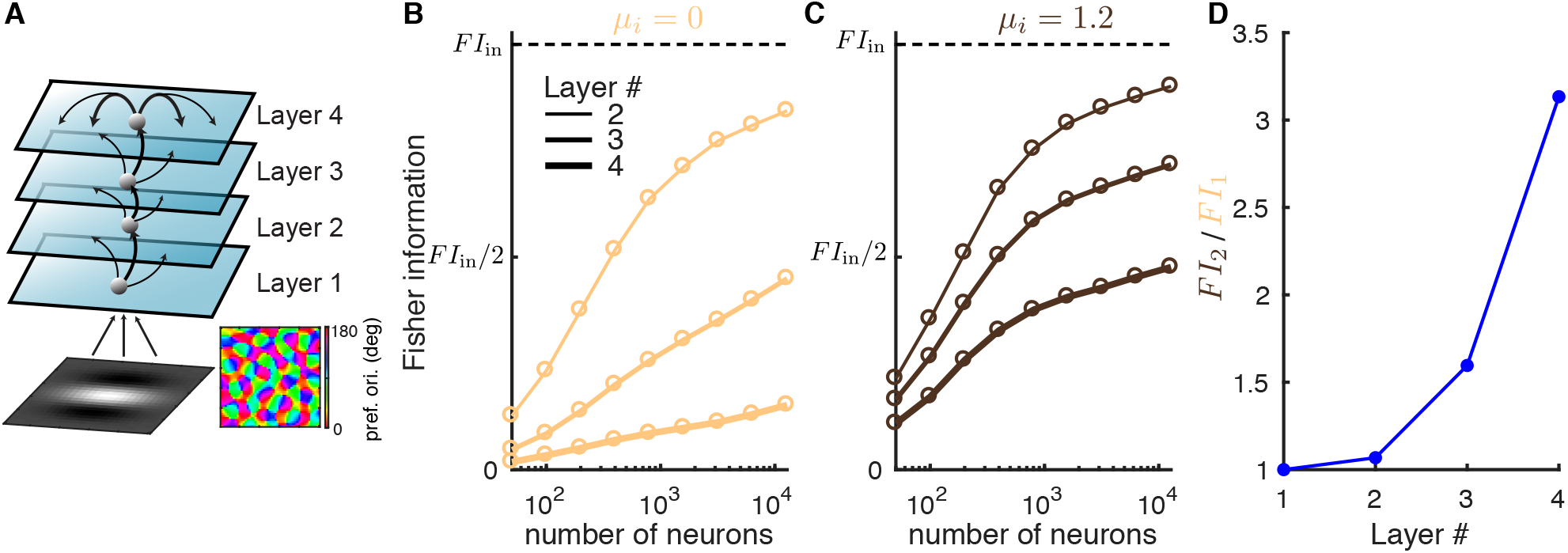
Information deteriorates in a multi-layer model with internal dynamics. **A,** Schematic of a multi-layer model. Layer 2-4 are recurrent networks of excitatory and inhibitory neurons, with the same projection widths of both recurrent and feedforward connections (*α*_rec_ = 0.1 and *α*_ffwd_ = 0.05). Recurrent connections of layer 2 and layer 3 are not shown for better illustration. Layer 1 network is the same as the V1 L4 network in Fig. 1A. **B,** The linear Fisher information is much more reduced in the higher-order layers when there is no input to the inhibitory neurons (*μ_i_* = 0). Dashed line is the total input Fisher information from layer 1 (*F I*_in_). **C,** Same as **B** with larger input to the inhibitory neurons (*μ_i_* = 1.2). D, The ratio between the Fisher information in networks with *μ_i_* = 1.2 (FI2) and that with *μ_i_* = 0 (FI1) across layers, encoded in a large neuron ensemble size (*N* = 12800). The feedforward strengths are adjusted such that the firing rates are similar between the two conditions of *μ_i_*, and are maintained across layers (Supp Fig. S1C).

## Discussion

The performance of a population code depends on the relationship between the individual tuning curves and the shared variability among neurons^22,26,27^. Previous studies have tacitly assumed a prescribed structure between tuning and variability, allowing a dissection of their respective impacts on population coding^23,31–33,81^. While this approach has given some critical insights, an understanding of how the mechanics of a neural circuit impact population coding has remained elusive. It is well known that the trial averaged^82,83^ and trial-to-trial variability^29,41,46,47^ of a population response are shaped by both feedforward and recurrent network connectivity. In this study we estimated the information available (to a linear decoder) about a simple, one dimensional stimulus that is contained in the activity of a recurrently coupled network of spatially ordered spiking neuron models. We show that simply understanding how pairwise noise correlations are determined through circuitry is insufficient to predict how information will be transferred across layers. Rather, an analysis of how circuitry determines the stability of network firing activity provides a better understanding of information transfer.

A central result of our paper is that in the limit of a large number of decoded neurons the linear Fisher information *I_F_* can either saturate to the input information 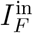 for networks with stable firing rate dynamics (Figs. 3,4), or fall short of that bound for networks in a pattern forming regime (Figs. 6,8,9). To appreciate why this is the case it is useful to recall a simple fact about how linear systems encode inputs. Consider a one dimensional stimulus *θ* that drives a vector of noisy inputs **x**(*θ*) = **s**(*θ*) + *η*, where the noise process *η* is zero mean with covariance matrix Σ_*x*_. Let the output **y**(*θ*) be a linear mapping of the input; specifically we take **y**(*θ*) = *L***x**(*θ*) for some invertible matrix *L*. Then we have that the linear Fisher information available to a decoder from **y**, termed 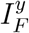, is simply:

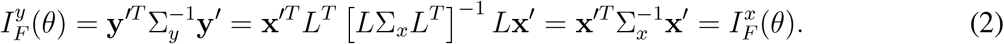

In other words, deterministic linear mappings do not distort or degrade information transfer (when the decoder is also linear). We remark that 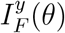 does not depend on the linear mapping *L*.

When the cortical network is in the stable regime and any input noise is weakly correlated then we can linearize network dynamics about an operating point^37,73,84,85^. In this regime the noisy spike counts during a trial, **n**(*θ*), in response to stimulus *θ* obey:

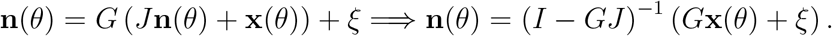

Here *G_ij_ = g_i_δ_ij_* is a diagonal matrix of the neuron response gains (depends on the operating point) and *J_ij_* is the synaptic coupling from neuron *j* to *i*. In addition to stimulus noise in **x**(*θ*) there is an internal noise process *ξ* that can reflect any trial fluctuations accrued from the digitization from continuous inputs to spike counts^35^ or a ‘background’ noise that is present even when coupling and stimulus are absent^37^; in all cases is independent across neurons. If sufficiently large numbers of neurons are decoded or sufficiently long observation times are used then the contribution of *ξ* to population statistics is negligible and we have that **n** ≈ (*I − GJ*)^−1^ *G***x**, or in other words the population output is a linear function of the input. Given this linearization and Eq. (2) we would then expect that 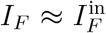, so long as the coupling *J* and input **x** provides a stable operating point around which to linearize. In contrast to this stable case, when the network is in a pattern forming regime then there is not a stable operating point about which to linearize. This means that Eq. (2) is no longer valid and we would expect that 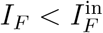. In total, this argument sketches how an understanding of the network firing rate stability translates quite naturally to an understanding of what circuit conditions will allow the network to faithfully transfer all the information available (to a linear decoder) in an input signal.

Broad inhibitory projections, also referred to as lateral inhibition, is a common circuit structure used to support various neural computations, such as working memory^71,86^, sharpening of tuning curves^29,82^, formation of clustered networks^87^, and the periodic spatial receptive fields of grid cells^88^. In this work, we show that the pattern-forming dynamics generated by lateral inhibition degrade information transmission, which is consistent with previous results^29^. Anatomical measurements of local cortical circuitry in visual cortex show that excitatory and inhibitory neurons project with similar spatial scales^89,90^. Our results suggest that such spatially balanced excitatory and inhibitory projections are important for maintaining faithful representations of sensory stimuli. Nevertheless, lateral inhibition may be important for transforming sensory information into task-related information in higher order cortices, such as prefrontal cortex.

Information transmission in feedforward networks has been studied in several models. Bejjanki and colleagues^91^ show that by improving feedforward template matching of Gabor filters, perceptual learning can increase the information gleaned from a presented image. In our model, the L4 network is similar to such models^41,91^, where information loss depends on the Gabor filters of the L4 neurons. In contrast, the feedforward projections from L4 to L2/3 neurons in our model are random and expansive (meaning there are much more neurons in L2/3 than L4), which is known to minimize information loss^30,92^. Therefore, there is little information loss in the feedforward projections from L4 to L2/3 in our model, regardless of the projection width. In addition, Renart and van Rossum^93^ and Zylberberg and colleagues^92^ study information transmission with external noise added to the output layer, and compute the optimal connectivity and input covariance, respectively, that maximize the information in the output layer. In our model, there is no external noise imposed on the L2/3 neurons. All the neuronal variability in L2/3 network is either internally generated or inherited from the feedforward inputs^42,46,47^.

Our results can potentially explain the improvement in discriminability by selective attention. Spatial attention has been shown to significantly reduce the global fluctuations in cortical recordings and meanwhile improve animals’ performance in orientation change detection task^15,16,94,95^. However, the magnitude of global fluctuations is found to have little effect on information, and instead information is only limited by input noise^35,41,81^. Recently, we developed a circuit model where large-scale wave dynamics give rise to low-dimensional shared variability^47^, thus capturing properties of population recordings in visual cortex^9,58,66,68,96^. By depolarizing the inhibitory neurons, the attentional modulation in our model stabilizes the network and decreases noise correlations. Therefore, the present study suggests that attention can improve information flow by quenching the internal turbulent dynamics in the recurrent circuit (Fig. 8A).

Spontaneous dynamics of cortex have been shown to mimic the response patterns evoked by external stimuli^97–99^. It has been hypothesized that the spontaneous dynamics reflect the prior distribution of the stimulus statistics, which is critical for optimal Bayesian inference^99,100^. Interestingly, our results show that the internal dynamics of recurrent networks do not improve information flow and can reduce it drastically when they are excessive. This is consistent with the correlation between the reduction in global fluctuations in population activity by attention and learning, and the improvement in animal’s performance^17,18^. However, the spontaneous dynamics in our model is not related to the stimulus since we only consider spatial wiring in the present study. Feature dependent wiring can generate spontaneous dynamics that resemble evoked responses^101^, and is likely to affect information transmission differently. The exact role of spontaneous dynamics in information processing as well as neural computation remains to be elucidated in future studies.

## Methods

### Network model description

The network consists of two stages, modeling for the layer 4 (L4) and layer 2/3 (L2/3) neurons in V1 respectively (Fig. 1A). Neurons on the two layers are arranged on a uniform grid covering a unit square Γ = [−0.5,0.5] × [−0.5,0.5]. L4 population is modeled as in Ref.^41^. There are *N_x_* = 2, 500 neurons in L4. The firing rate of each neuron is

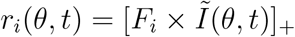

where 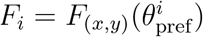 is a Gabor filter and 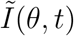 is a Gabor image corrupted by independent noise following Ornstein-Uhlenbeck process.

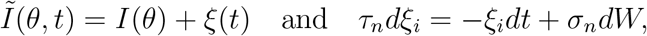

with *τ_n_* = 40 ms and *σ_n_* = 3.5. The orientation *θ* is normalized between 0 and 1. Spike trains of L4 neurons are generated as inhomogeneous Poisson process with rate *r_i_*(*t*).

The Gabor image is defined on Γ with 25 × 25 pixels (Δ*x* = 0.04),

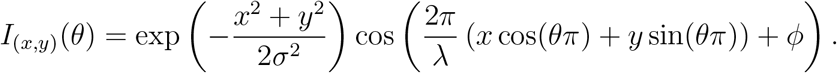

The size of the spatial Gaussian envelope was *σ* = 0.2, and the spatial wavelength was *λ* = 0.6 and phase was *φ* = 0. The Gabor filters were

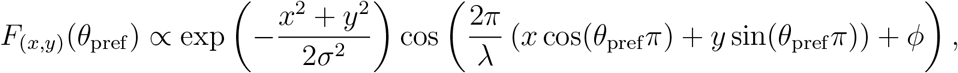

with the same *σ, λ* and *φ* as the image. The orientation map was generated using the formula from Ref.^53^ (Supp Materials Eq. 20). The preferred orientation at (*x,y*)is *θ*_pref_ (*x, y*) = angle(*z*(*x,y*))/(2*π*) and

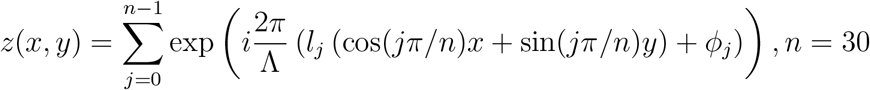

where Λ = 0.2 is the average column spacing, *l_j_* = ±1 is a random binary vector and the phase *φ_j_* is uniformly distributed in [0, 2*π*].

The L2/3 network consists of recurrently coupled excitatory (*N*_e_ = 40, 000) and inhibitory (*N*_i_ = 10, 000) neurons. Each neuron is modeled as an exponential integrate-and-fire (EIF) neuron whose membrane potential is described by:

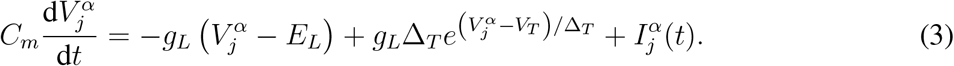

Each time 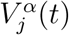 exceeds a threshold *V*_th_, the neuron spikes and the membrane potential is held for a refractory period *τ*_ref_ then reset to a fixed value *V*_re_. Neuron parameters for excitatory neurons are *τ*_m_ = *C_m_/g_L_* = 15 ms, *E_L_* = −60 mV, *V_T_* = −50 mV, *V*_th_ = −10 mV, Δ_*T*_ = 2 mV, *V*_re_ = −65 mV and *τ*_ref_ = 1.5 ms. Inhibitory neurons are the same except *τ_m_* = 10 ms, Δ_*T*_ = 0.5 mV and *τ*_ref_ = 0.5 ms. The total current to each neuron is:

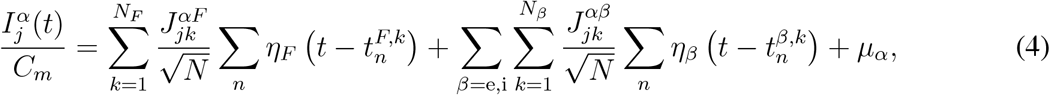

where *N = N_e_ + N_i_* is the total number of neurons in the network. Postsynaptic current is

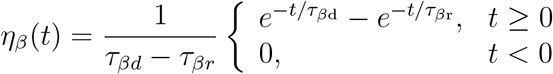

where *τ*_er_ = 1 ms, *τ*_ed_ = 5 ms for excitatory synapses and *τ*_ir_ = 1 ms, *τ*_id_ = 8 ms for inhibitory synapses. The feedforward synapses from L4 to L2/3 have the same kinetics as the recurrent excitatory synapse, i.e. *η_F_*(*t*) = *η*_e_(*t*).

The probability that two neurons, with coordinates **x** = (*x*_1_, *x*_2_) and **y** = (*y*_1_, *y*_2_) respectively, are connected depends on their distance measured on Γ with periodic boundary condition:

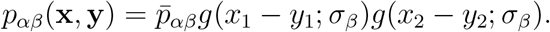

Here 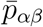 is the mean connection probability and

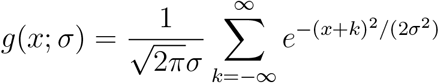

is a wrapped Gaussian distribution. A presynaptic neuron is allowed to make more than one synaptic connection to a single postsynaptic neuron.

The mean recurrent connection probabilities were 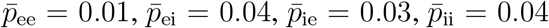 and the recurrent synaptic weights were *J*_ee_ = 80 mV, *J*_ei_ = −240 mV, *J*_ie_ = 40 mV and *J*_ii_ = −300 mV. The feedforward connection probabilities were 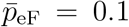 and 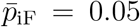. The feedforward connection strengths were *J*_eF_ = 240 mV and *J*_iF_ = 400 mV for Figure 1-6, and *J*_eF_ = 140 mV and *J*_iF_ = 0 mV for Figure 7 and 8. The static inputs to inhibitory neurons were *μ_i_* = 0 mV/ms for Figure 1-6, and *μ_i_* = 0, 0.4, 0.8, 1.2 mV/ms for Figure 7 and 8. The static inputs to excitatory neurons were *μ_e_* = 0 mV/ms for all simulations.

The feedforward connection widths, *α*_ffwd_, were 0.05, 0.625, 0.75, 0.1 and ∞ and the excitatory and inhibitory connection widths were fixed to be *σ*_e_ = *σ*_i_ = 2*α*_ffwd_ for Figure 2 and 3. The inhibitory connection widths *σ_i_* were 0.1, 0.2 and 0.3 for Figure 5 and 6, and the excitatory and the feedforward connection widths were *σ*_e_ = 0.1 and *α*_ffwd_ = 0.05 respectively. The feedforward connection width was *α*_ffwd_ = 0.05 and the excitatory and inhibitory connection widths were *σ*_e_ = *σ*_i_ = 0.1 for Figure 1,7-9.

In the multi-layer network (Fig 9), the recurrent and the feedforward connections of all layers are *α*_rec_ = 0.1 and *α*_ffwd_ = 0.05, respectively. The first and the second layer are the same as the L4 and L2/3 networks, respectively, described above. The other layers are modeled the same as the L2/3 network. In the condition when *μ_i_* = 0 mV/ms, the feedforward connection strength from layer 1 to layer 2 excitatory neurons is *J*_eF_ = 140 mV, and the feedforward connection strengths between other layers are *J*_eF_ = 35 mV. In the condition when *μ_i_* = 1.2 mV/ms, the feedforward connection strength from layer 1 to layer 2 excitatory neurons is *J*_eF_ = 180 mV, and the feedforward connection strengths between other layers are *J*_eF_ = 54 mV. All the feedforward connections to inhibitory neurons and the static inputs to excitatory neurons were set to be zero, i.e. *J*_iF_ = 0 mV and *μ_e_* = 0 mV/ms. The feedforward strengths were chosen such that the mean firing rates of layer 2-4 are similar across layers as well as between the two conditions of *μ_i_*.

All simulations were performed on the CNBC Cluster in the University of Pittsburgh. All simulations were written in a combination of C and Matlab (Matlab R 2015a, Mathworks). The differential equations of the neuron model were solved using forward Euler method with time step 0.05 ms.

### Neural field model and stability analysis

We use a two dimensional neural field model to describe the dynamics of population rate (Fig. 4 and 6). The neural field equations are

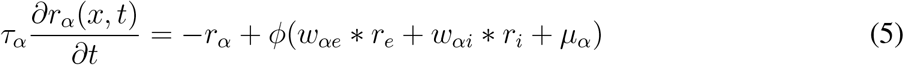

where *r_α_*(*x, t*) is the firing rate of neurons in population *α* = e, i near spatial coordinates *x* ∈ [0,1] × [0, 1]. The symbol ∗ denotes convolution in space, *μ_α_* is a constant external input and 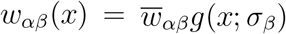 where *g*(*x; σ_β_*) is a two-dimensional wrapped Gaussian with width parameter *σ_β_, β* = e, i. The transfer function is a threshold-quadratic function, 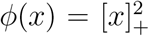. The timescale of synaptic and firing rate responses are implicitly combined into *τ_α_*.

For constant inputs, *μ_e_* and *μ_i_*, there exists a spatially uniform fixed point, 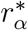. Linearizing around this fixed point in Fourier domain gives a Jacobian matrix at each spatial Fourier mode^45^

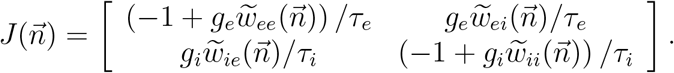

where 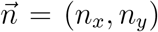 is the two-dimensional Fourier mode, 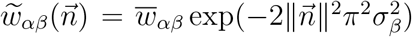 is the Fourier coefficient of *w_αβ_*(*x*) with 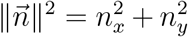 and 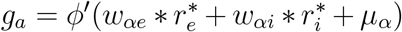 is the gain. The fixed point is stable at Fourier mode 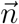 if both eigenvalues of 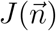 have negative real part. Note that stability only depends on the wave number, 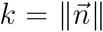, so Turing-Hopf instabilities always occur simultaneously at all Fourier modes with the same wave number (spatial frequency).

For the stability analysis in Figures 5 and 7, *μ_i_* varied from 0.1 to 0.7 with step size 0.002, *σ_i_* varied from 0.05 to 0.2 with step size 0.0005, and *μ*_e_ = 0.5 and *σ*_e_ = 0.1. The rest of the parameters were 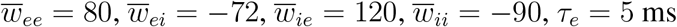 and *τ_i_* = 8 ms.

### Statistical methods

Each simulation was 20 sec long consisting of alternating OFF (300 ms) and ON (200 ms) intervals. Image was presented during ON intervals, where the average firing rate of L4 neurons was *r_X_* = 10 Hz. During OFF intervals, spike trains of L4 neurons were independent Poisson process with rate *r_X_* = 5 Hz. Spike counts from the L2/3 excitatory neurons during the ON intervals were used to compute the linear Fisher information and noise correlations. The first spike count in each simulation was excluded. For each parameter condition, the connectivity matrices were fixed for all simulations. The initial states of each neuron’s membrane potential were randomized in each simulation.

To compute tuning curve functions, the orientation for each ON interval was randomly sampled from 50 orientations uniformly spaced between 0 and 1. There were 9,750 spike counts in total for all orientations. Tuning curves were smoothed with a Gaussian kernel of width 0.05. Orientation selectivity index for neuron *i* is computed as

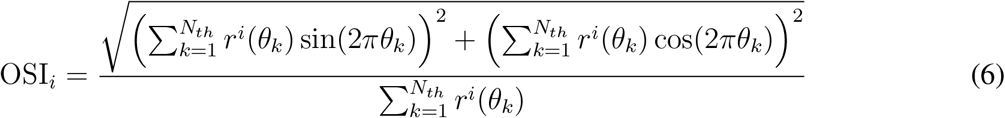

where *r^i^*(*θ*) is the tuning curve function of neuron *i*, and *θ_k_* = *k/N*_th_.

To compute the linear Fisher information and noise correlation, the orientations of the Gabor images during ON intervals, were randomly chosen from *θ*_1_ = *θ + δθ*/2 and *θ*_2_ = *θ - δθ*/2, where *θ* = 0.5 and *δθ* = 0.01. The linear Fisher information of L2/3 neurons is computed using the bias-corrected estimate^64^,

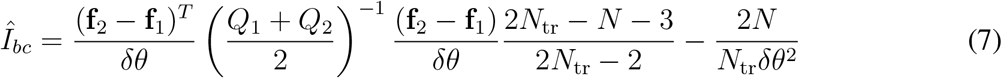

where **f**_*i*_ and *Q_i_* are the empirical mean and covariance, respectively, for *θ_i_*. *N*_tr_ was the number of trials for each *θ_i_*. The number of neurons were *N* = 50, 100, 200, 400, 800, 1600, 3200, 6400, 12800, randomly sampled without replacement from the excitatory population of L2/3.

The linear Fisher information of L4 neurons can be estimated analytically, where

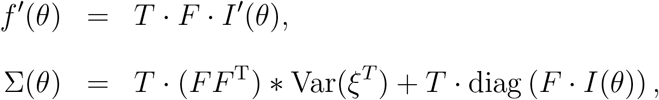

with 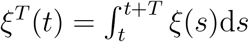 and *T* be the time window for spike counts. With *T* = 200 ms and *τ_n_* = 40 ms, 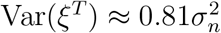.

Th noise correlation was computed with *N* = 1600 neurons randomly sampled without replacement from the excitatory population of L2/3. The Pearson correlation coefficients were computed from the average covariance matrix (*Q*_1_ + *Q*_2_)/2.

For the computation of both noise correlations and linear Fisher information, there were 10 sampling of neurons for each *N*. Neurons of firing rates less than 1 Hz were excluded. There were 58,500 spike counts in total for *θ*_1_ and *θ*_2_.

### Code availability

Computer code for all simulations and analysis of the resulting data will be available at https://github.com/hcc11/.

## Acknowledgments

The work was funded by the Swartz Foundation Fellowship #2017-7 (C.H.), grants from the Simons Foundation Collaboration on the Global Brain (C.H., A.P and B.D.), the Swiss National Science Foundation (www.snf.ch), #31003A_143707 and #31003A_165831 (A.P.), National Institutes of Health (Grants #1U19NS107613-01 and #R01 EB026953), and the Vannevar Bush Faculty (Fellowship #N00014-18-1-2002). This work used the Extreme Science and Engineering Discovery Environment (XSEDE), which is supported by National Science Foundation grant number ACI-1548562. Specifically, it used the Bridges system, which is supported by NSF award number ACI-1445606, at the Pittsburgh Supercomputing Center (PSC).

## Author Contributions

C.H., A.P., and B.D. conceived the project; C.H. performed the simulations and data analysis; B.D. supervised the project; all authors contributed to writing the manuscript.

## Author Information

The authors declare no competing financial interests. Correspondence and requests for materials should be addressed to B.D.

