## Supplemental Information for "Internally generated population activity in cortical networks hinders information transmission"

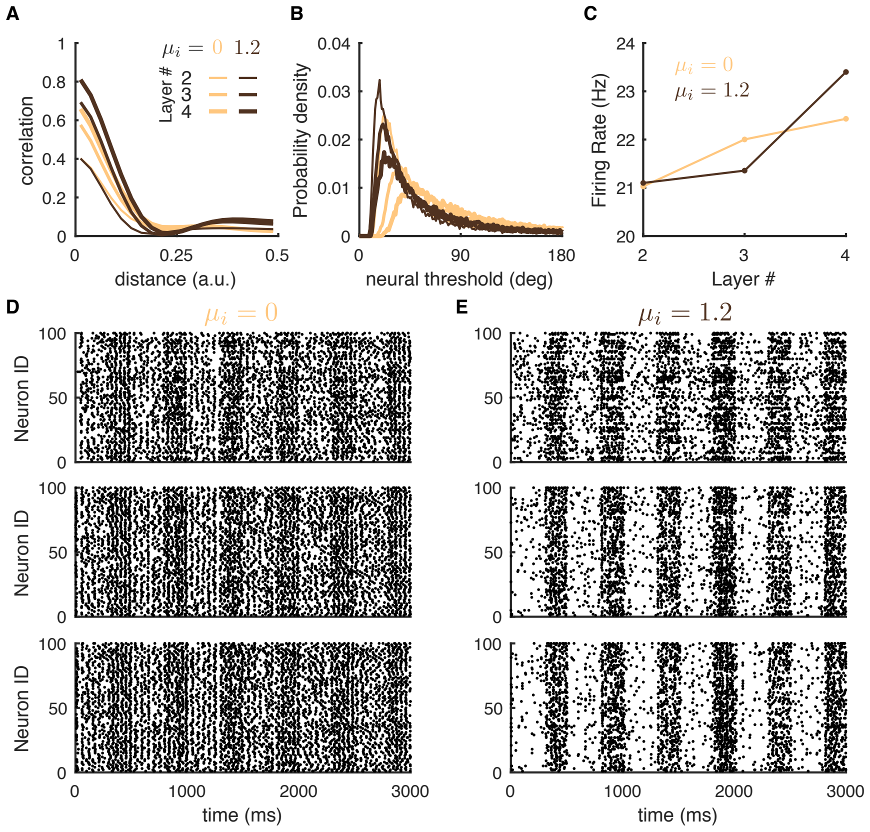


Figure S1: **Related to Figure 9.** The network responses in a multi-layer model with (orangel, $\mu_i = 0$) or without (dark brown, $\mu_i = 1.2$) depolarizing currents to the inhibitory neurons ($\mu_i$) across layers. **A**, Pairwise correlations as a function of the distance between excitatory neuron pairs across layers (Layer 2, thin lines; Layer 3; medium-width lines; Layer 4; thick lines). **B**, The probability density distributions of neural thresholds ($\sigma_i/f_i'$) of excitatory neurons across layers and in the two conditions of $\mu_i$. **C**, The mean fring rates of the excitatory populations across layers. The feedforward strengths between layers were chosen such that the mean firing rates of layer 2-4 were similar across layers as well as between the two conditions of $\mu_i$. **D**, The raster plots of 100 excitatory neurons randomly selected from Layer 2 to Layer 4 (top to bottom panel) in a network without depolarizing current to the inhibitory neurons ($\mu_i = 0$). The image was presented for 200 ms and then off for 300 ms in each trial. The average rates of Layer 1 neurons were 10 Hz during image presentation and 5 Hz when image was off. **E**, Same as **D** for a network with $\mu_i = 1.2$. The inputs from Layer 1 are the same in both conditions of $\mu_i$. See Methods for model details.
